# Opto-p53: A Light-Controllable p53 Signaling for Regulating p53-Dependent Cell Fate

**DOI:** 10.1101/2025.04.04.647217

**Authors:** Tatsuki Tsuruoka, Yuhei Goto, Kazuhiro Aoki

## Abstract

p53 protein, a crucial transcription factor in cellular responses to a wide variety of stress, regulates multiple target genes involved in tumor suppression, senescence induction, and metabolic functions. However, it remains unclear how diverse cellular phenotypes are modulated by p53. In this study, we developed an optogenetic tool, Opto-p53, to control p53 signaling by light. Opto-p53 was designed to trigger p53 signaling by reconstituting p53 N-terminal and C-terminal fragments with a light-inducible dimerization (LID) system. Upon light exposure, cells expressing Opto-p53 demonstrated p53 transcriptional activation, resulting in cell death and cell cycle arrest. We further enhanced the efficacy of light-induced p53 activation by introducing specific mutations into Opto-p53 fragments. Our findings unveil the capability of Opto-p53 to serve as a powerful tool for dissecting the complex roles of p53 in cellular processes, thereby contributing to the field of synthetic biology and providing general design principles for optogenetic tools using endogenous transcription factors.

## Introduction

The tumor suppressor protein p53 is known to function as a hub transcription factor for various stress signals and to regulate the expression of hundreds of target genes (Fischer, 2017). In recent years, it has become clear that p53 functions not only in tumor suppression but also in a wide variety of other areas, such as senescence induction, metabolic regulation, and stem cell differentiation (Jain and Barton, 2018; Labuschagne et al., 2018; Sheekey and Narita, 2023). The question of how a single signaling factor can induce diverse cellular phenotypes has attracted the attention of many researchers.

Classically, the idea has been proposed that p53 activates distinct gene sets through post-translational modification (PTM) patterns and interacting factors that depend on upstream inputs, resulting in a variety of phenotypic outputs (Murray-Zmijewski et al., 2008). However, recent studies have shown that the different temporal dynamics of p53 activation can also yield distinct cellular phenotypes (Batchelor and Loewer, 2017). Given the multifaceted roles of p53, synthetic biology approaches that can activate the target factors in a context-dependent manner are a promising strategy for functional analysis.

Optogenetics is a type of synthetic biology that has been used more and more in different research areas (Repina et al., 2017). This is because optogenetics allows the manipulation of various biological processes by light with high spatial and temporal resolutions. Most optogenetic tools are derived from photo-responsive proteins found in plants, fungi, and bacteria. This allows scientists to precisely and bio-orthogonally perturb various intracellular events such as opening ion channels, regulating protein-protein interaction, and cell signaling (Lan *et al*., 2022; Manoilov et al., 2021).

One of the most successful applications of optogenetics is the regulation of gene expression by light. In research using mammalian cells, several synthetic transcription factors have been reported that combine DNA-binding domains (DBD) and transcription activation domains (TAD) with a light-induced dimerization (LID) system (Guntas *et al*., 2015; Kawano *et al*., 2015; Kennedy et al., 2010; Levskaya et al., 2009; Yazawa et al., 2009; Zoltowski and Crane, 2008). Some of the most commonly used DBDs and TADs come from Gal4 or TetR and VP16 or NF-κB, respectively (Kennedy et al., 2010; Levskaya et al., 2009; Schneider *et al*., 2021; Wang *et al*., 2012; Yamada et al., 2018; Yazawa *et* al., 2009). These light-responsive transcription factors are highly bio-orthogonal, thereby selectively inducing gene expression from exogenous gene cassettes in response to light. There have also been attempts to regulate endogenous gene expression. For this purpose, dead Cas9 (dCas9)-based systems have been widely employed as a DBD to recruit synthetic transcription factors to the specific endogenous gene locus (Gao et al., 2016; Kim et al., 2023; Nihongaki *et al*., 2015; Polstein and Gersbach, 2015). One advantage of these systems is that the target gene can be easily altered by changing the sequence of the sgRNA. However, the aforementioned optogenetic tools allow, in principle, to regulate the expression of only one or a few genes. Indeed, there are a number of endogenous transcription factors that function as a hub linking multiple upstream signaling and the transcriptional regulation of diverse target genes (Lambert et al., 2018). To mimic the activation of multiple target genes regulated by such transcription factors, one possible strategy is to utilize the DBD and TAD derived from endogenous transcription factors.

In this study, we present optogenetic tools for reconstituting the DBD and TAD of p53, Opto-p53, to trigger gene expression and downstream phenotypes of p53, such as cell death and cell cycle arrest. This new platform of Opto-p53 has the potential to be applied to the light-dependent manipulation of various other transcription factors.

## Results

### Design of the Opto-p53 system

We aim to recapitulate p53 signaling by light and examine how p53 activation induces gene expression and phenotype. For this purpose, we attempt to design an optogenetic system allowing light-dependent reconstitution of functional p53 in living cells. p53, a 393 amino acid (a.a.) protein, consists of five domains including a transcriptional activation domain (TAD) and a DNA binding domain (DBD) (Fig. 1A). Hence, first, we split p53 into two fragments: p53 N-terminus fragment (p53NT, 1–97 a.a.) containing the TAD and p53 C-terminus fragment (p53CT, 98–393 a.a.) containing the DBD. Of note, p53NT contains S15D phospho-mimetic mutation, which mimics p53 activation by upstream kinases (Sakaguchi *et al*., 1998). Second, we introduce a light-inducible dimerization (LID) system for reconstituting p53 by bringing p53NT and p53CT in close proximity to each other (Fig. 1B). As a LID tool, a blue light-responsive protein cryptochrome 2 (CRY2) and its binding domain CIBN are adopted because of the high reproducibility (Kennedy et al., 2010). This optogenetic system is hereafter referred to as Opto-p53. The DBD-containing fragment and the TAD-containing fragment are designated as the localizer and the actuator, respectively (Fig. 1B). Each fragment is fused to a different fluorescent protein to confirm its expression and subcellular localization. A nuclear localization signal (NLS) is also attached to the actuator fragment to enhance its nuclear localization. Previous studies have shown that the DBD of p53 binds to p53-responsive elements in the genome with high affinity (Krois *et al*., 2018; Weinberg *et al*., 2005; Weinberg, Veprintsev, et al., 2004). Therefore, p53CT is expected to be constitutively associated with p53-responsive elements in the promoter region of the target gene. Blue light illumination causes the interaction between CRY2 and CIBN, leading to the recruitment of p53NT to p53CT located at the target gene promoter. This, in turn, triggers the recruitment of transcription coactivators including p300/CBP, and induces p53-dependent transcriptional activation (Fig. 1C).

**Figure 1.**
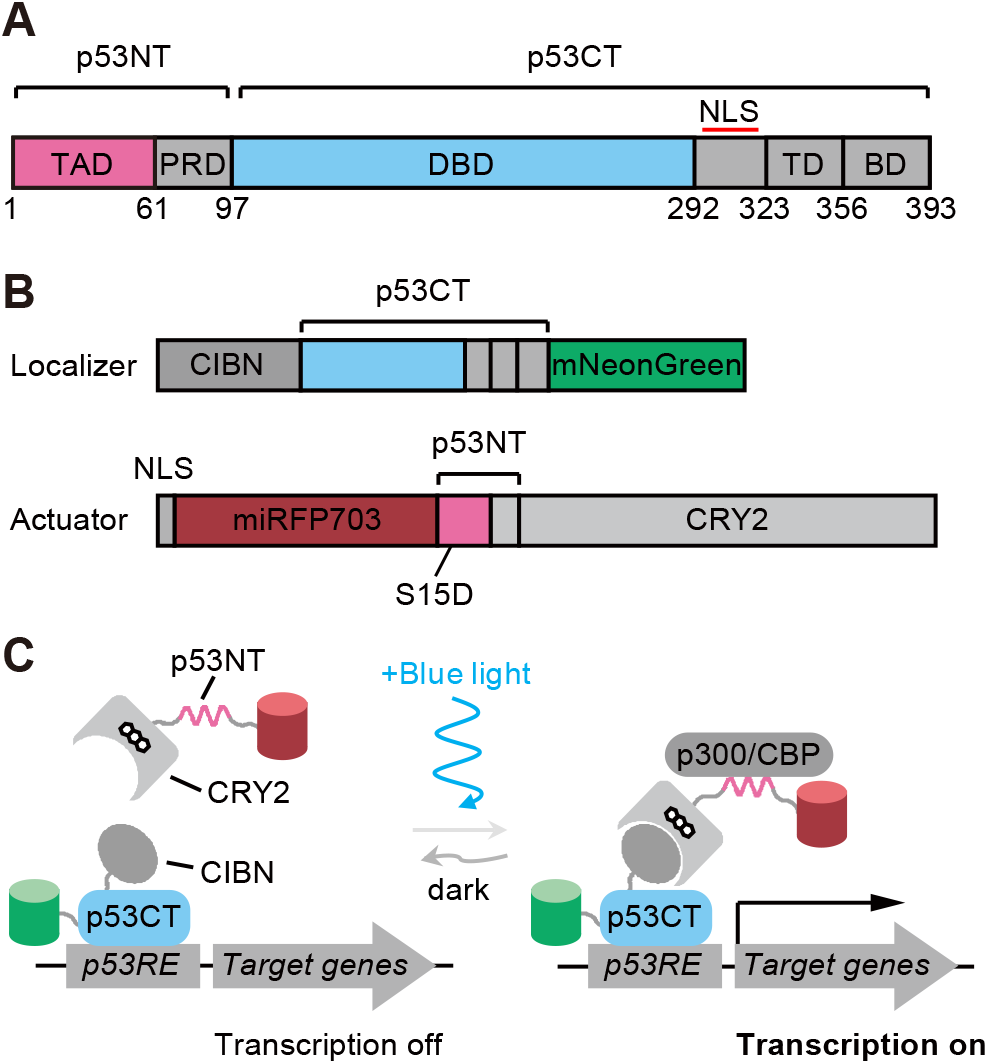
Design of the Opto-p53 system. A. Domain structure of the p53. TAD, Transactivatioan domain; PRD, Proline-rich domain; DBD, DNA binding domain; TD, Tetramerization domain; BD, Basic domain; NLS, Nuclear localization signal; p53NT, p53 N-terminal fragment; p53CT, p53 C-terminal fragment. B. Expression constructs of the Opto-p53 system. Upper panel: p53CT localizer. Lower panel: p53NT actuator. p53NT contains a phospho-mimetic mutation, S15D, which mimics p53 activation by upstream kinases. C. Schematic illustration of the Opto-p53 system. *p53RE*: p53 responsive element, which is located in the promoter region of the p53 target gene.

### Characterization of Opto-p53 fragments expressed in living cells

CRY2 is known to possess hetero-dimerization activity with CIBN and homo-oligomerization activity, which gives rise to droplet-like structures through liquid-liquid phase separation (LLPS)(Bugaj *et al*., 2013; Wang and Lin, 2025). Several studies have reported that forming such LLPS condensate promotes the transcriptional activation of synthetic transcription factors (Hnisz et al., 2017; Schneider et al., 2021; Wei *et al*., 2020; Wu *et al*., 2022). Therefore, we examined the subcellular localization of Opto-p53 fragments under blue light illumination.

When only the p53CT localizer fragment containing CIBN was expressed in HCT116 cells, it showed the same nuclear localization as a nuclear marker Histone H1-mCherry, and no light-dependent changes in the localization under blue light exposure (Fig. 2A, upper). Meanwhile, the CRY2-containing p53NT actuator fragment, which was uniformly localized to the nucleus under the dark condition, formed small foci in the nucleus in a blue light-dependent manner (Movie S1) (Fig. 2A, middle). The formation of these foci was reversible and could be repeatedly induced by switching blue light on and off. A similar light-dependent localization change was observed in the actuator fragment with three tandem minimal transactivation domains derived from VP16 (VP16minADx3) instead of p53NT, which was prepared as a positive control for TAD of synthetic transcription factors (Fig. 2A, lower) (Seipel *et al*., 1992). To quantify the localization changes in the actuator fragments, we analyzed the time course of the coefficient of variation (CV) of the miRFP703 signal in each nucleus (Fig. 2B). CV value is calculated by dividing the standard deviation by the average value, thereby representing to what extent the fluorescent intensity in the nucleus is uneven. We found that the light-induced change in CV was slightly higher in the actuator with p53NT than in VP16minADx3. Given that intrinsically disordered regions (IDRs) generally promote droplet formation through LLPS, the IDR nature of p53NT may contribute to this difference (Dyson and Wright, 2025). Meanwhile, CRY2 proteins were overexpressed, and foci were only observed in cells with high expression levels (Fig. S1A). As a result, the change in CV value was small (Fig. S1B).

**Figure 2.**
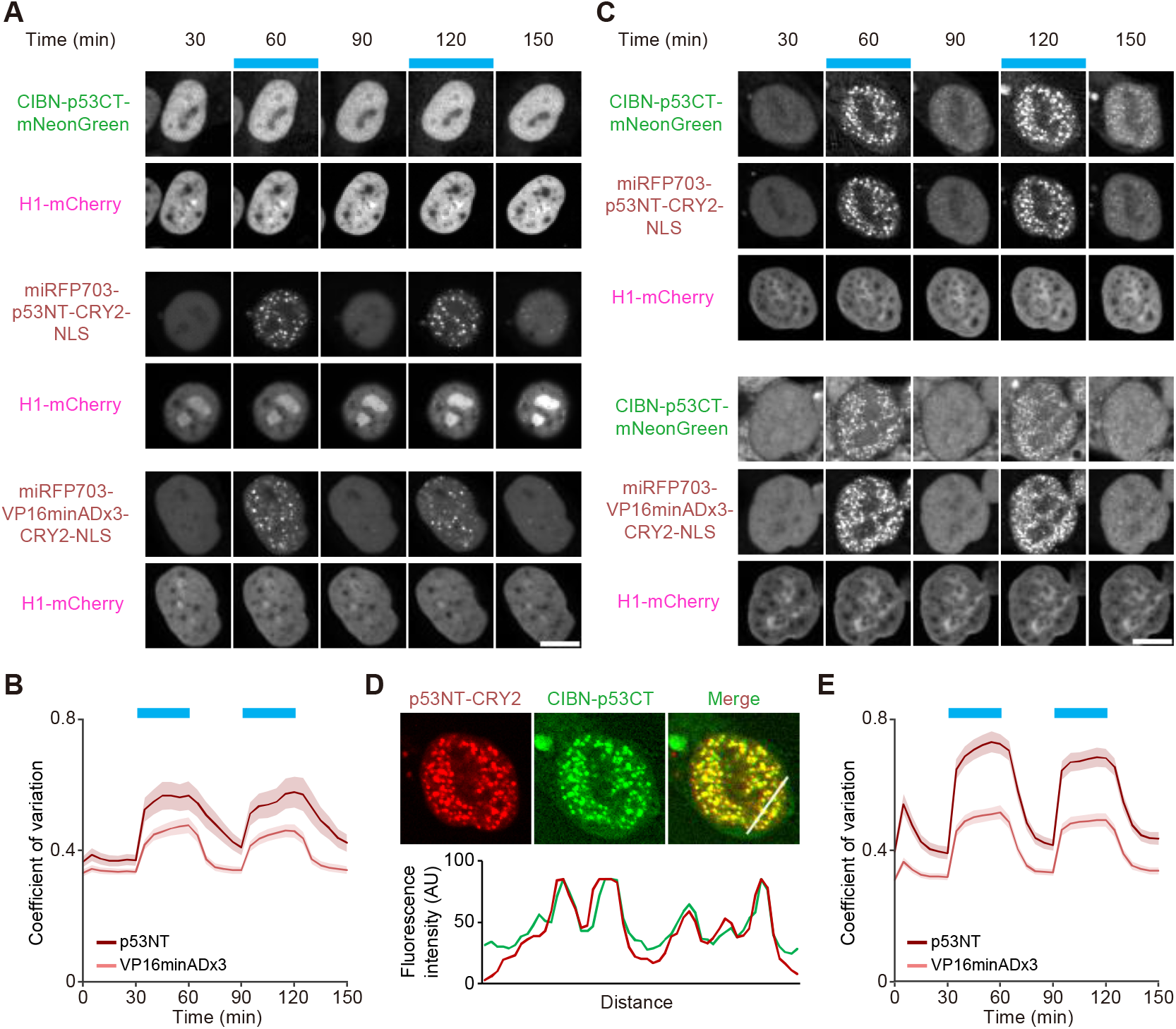
Light-dependent changes in the localization of the Opto-p53 actuator and localizer. A. Light-dependent changes in subcellular localization when the Opto-p53 actuator or localizer is expressed alone in HCT116 cells. The blue boxes indicate the time points at which blue light illumination was applied. H1-mCherry is a nuclear marker. Scale bar, 10 μm. B. Temporal changes in the coefficient of variation of nuclear miRFP703 fluorescence intensity in cells expressing Opto-p53 actuator fragments. The plot shows the mean ± s.e.m. p53NT, n = 35 cells; VP16minADx3, n = 39 cells. C. Light-dependent changes in subcellular localization when the Opto-p53 actuator and localizer were co-expressed. Scale bar, 10 μm. D. Co-localization of the Opto-p53 actuator and localizer by blue light illumination. Upper panel: Representative images of the Opto-p53 actuator, localizer, and their merged image. The cell in these images was the same as in the upper panels of Figure 2C. Lower panel: Fluorescence intensities along the white line shown in the merged image. E. Temporal changes in the coefficient of variation of nuclear miRFP703 fluorescence intensity in cells co-expressing Opto-p53 actuator and localizer. The plot shows the mean ± s.e.m. p53NT, n = 60 cells; VP16minADx3, n = 68 cells.

Next, we co-expressed both the localizer and the actuator of Opto-p53 in the cells. While both the p53NT actuator and the p53CT localizer were diffusely localized at the nucleus under the dark condition, not only the actuator but also the localizer formed bright foci at the nucleus in a light-dependent manner (Fig. 2C, upper). The line-scan profile indicates the co-localization of these foci in the nucleus (Fig. 2D). Similarly, the p53CT localizer also formed foci upon blue light exposure when VP16minADx3 was used as the TAD (Fig. 2C, lower). The time course of the CV values demonstrated light-induced foci formation of the p53NT actuator and VP16minADx3 with p53CT localizer in the nucleus (Fig. 2E). Interestingly, in the case of both p53NT and VP16minADx3, the increase in the CV values upon blue light exposure was larger under the co-expression condition (Fig. 2E) than that under the condition where only the actuator was expressed (Fig. 2B). This result could be due to the ability of tetramer formation through the tetramerization domain in the p53CT localizer, thereby enhancing the formation of CRY2-induced foci.

### Light-dependent transcriptional activation of the p53 pathway

We next asked whether Opto-p53 could induce transcriptional activation of p53 by light. To this end, we established a cell line stably harboring a p53 transcriptional reporter in the genome, which includes a p53 responsive element (*p53RE*) derived from *CDKN1A* promoter region followed by CMV minimal promoter and mScarlet-I-3xNLS (Fig. S2A) (Tsuruoka et al., 2023). When p53 is activated in cells harboring this p53 transcriptional reporter, p53 binds to the *p53RE* and induces the expression of mScarlet-I-3xNLS. Therefore, we can estimate the activity of p53 from the fluorescence intensity of mScarlet-I. Indeed, the treatment of these cells for 24 hours with etoposide, which is known to activate the p53 signaling by inducing DNA damage (Nitiss, 2009; Yang et al., 2018), increased mScarlet-I intensity in a dose-dependent manner (Fig. S2B). Similar results were also observed with nutlin-3a treatment, which selectively activates the p53 signaling by inhibiting the interaction of p53 with its negative regulators, MDM2/MDMX (Vassilev et al., 2004) (Fig. S2C). There was a much more significant increase in mScarlet-I intensity in the nutlin-3a-treated condition compared to the etoposide-treated condition, with a more than 10-fold increase at a drug concentration of 20 μM.

We transiently introduced the Opto-p53 fragments by lipofection into HCT116 cells harboring the p53 transcriptional reporter, and illuminated the cells with blue light for 24 hours. As expected, the cells expressing the p53NT actuator and the p53CT localizer gradually increased the mScarlet-I fluorescence intensity (Movie S2) (Fig. 3A, first row, and Fig. 3B). Furthermore, the combination of the p53CT localizer with the VP16minADx3 actuator also shows an increase in the p53 transcriptional reporter (Fig. 3A, second row, and Fig. 3B). On the other hand, neither the single expression of the actuator nor the localizer caused p53 transcriptional activation under the blue light conditions (Fig. 3A, third to fifth rows, and Fig. 3B). These results demonstrate that condensate formation of the TAD-containing actuator in the nucleus (Fig. 2A) does not suffice to induce p53 transcriptional activation, and that proper recruitment of the actuator to the promoter region through binding to the localizer leads to downstream gene expression. To our surprise, there was no substantial difference in p53 transcriptional activation when p53NT and VP16minADx3 were used as TAD (Fig. 3B). These data suggest that the p53NT actuator fragment is an efficient transcriptional activator comparable to the three tandem repeats of the VP16 minimal transcriptional activation domain, which have been widely used as a very potent transcriptional activation domain. We also confirmed that neither the single expression of p53NT actuator or p53CT localizer nor their co-expression caused p53 transcriptional activation under the dark condition (Fig. S2D and S2E).

**Figure 3.**
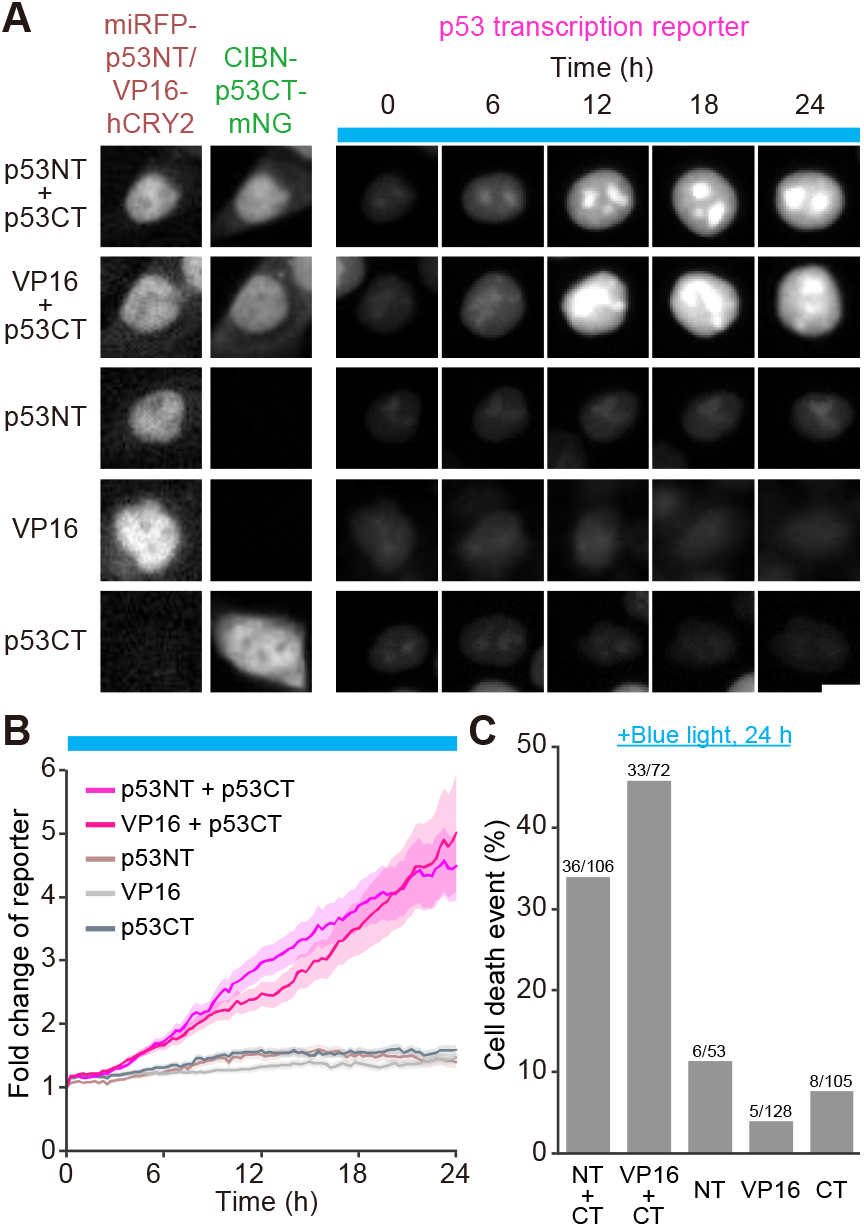
Transcriptional activation of the p53 by the blue light illumination. A. Expression patterns of Opto-p53 actuator or localizer and light-dependent changes in p53 transcription reporter. The blue boxes indicate the time points at which blue light illumination was applied. Scale bar, 10 μm. B. Fold changes in the p53 transcription reporter in each condition. The plot shows the mean ± s.e.m. p53NT + p53CT, n = 106 cells; VP16minADx3 + p53CT, n = 72 cells; p53NT, n = 53 cells; VP16minADx3, n = 128 cells; p53CT, n = 105 cells. C. Quantification of cell death in Figure 3B. The numbers on each grey bar indicate the number of dead cells/total cells.

Next, we investigated whether optogenetic activation of p53 causes phenotypic changes, and analyzed cell death because cell death is a major outcome of p53 activation (Riley *et al*., 2008). Dead cells were manually detected based on the morphological change of each nucleus (Fig. S2F). Consistent with the increase in transcriptional activity of p53, the percentage of cell death enhanced upon blue light exposure when the cells were co-expressing the p53CT localizer and the p53NT or VP16minADx3 actuator (Fig. 3C, first and second columns), indicating light-induced cell death through p53 activation. We note that a certain number of cell deaths was observed even under the single expression of these fragments (Fig. 3C, third to fifth columns). To investigate whether these cell deaths were due to the phototoxicity of blue light illumination, we cultured the cells expressing Opto-p53 fragments under dark conditions and found comparable levels of cell death (Fig. S2G). These results indicated that the basal cell death in Figure 3C is not due to the phototoxicity of blue light but possibly due to spontaneous cell death and/or cytotoxicity caused by transient overexpression.

### Cell cycle regulation by light-dependent p53 activation

As cell cycle arrest is also a major downstream of p53 activation, we finally tried to recapitulate the cell cycle arrest via p53 activation by blue light illumination using stable cell lines expressing Opto- p53. Because the expression level in stable cell lines is generally lower compared to transient expression, we modified the p53CT localizer and p53NT actuator fragments of Opto-p53 to enable more efficient p53 activation as follows. First, five phosphorylation-mimetic mutations were introduced into p53NT (p53NT(5SD)) to enhance the interaction between p53 and the transcriptional coactivator p300/CBP and to attenuate the interaction with the negative regulators MDM2 and MDMX (Jabbur and Zhang, 2002; Lee et al., 2010; Teufel et al., 2009)(Fig. 4A, upper). Similarly, six acetylation-mimetic mutations, which are known to stabilize p53 and increase transcriptional activity, were introduced into p53CT (p53CT(6KQ))(Fig. 4A, lower) (Barlev *et al*., 2001; Gu and Roeder, 1997; Kon et al., 2021; Luo et al., 2004; Wang *et al*., 2016; Weinberg, Freund, *et al*., 2004). As a negative control, we prepared p53NT with the mutation L22Q/W23S/W53Q/F54S, which disrupts binding to p300/CBP (p53NT(NC)), and p53CT with the mutation R273H in the DNA binding domain, which abolishes DNA binding (p53CT(R273H))(Fig. 4A) (Candau et al., 1997; Cho *et al*., 1994; Joerger *et* al., 2005; Lin et al., 1994; Teufel et al., 2007; Zhu et al., 1998).

**Figure 4.**
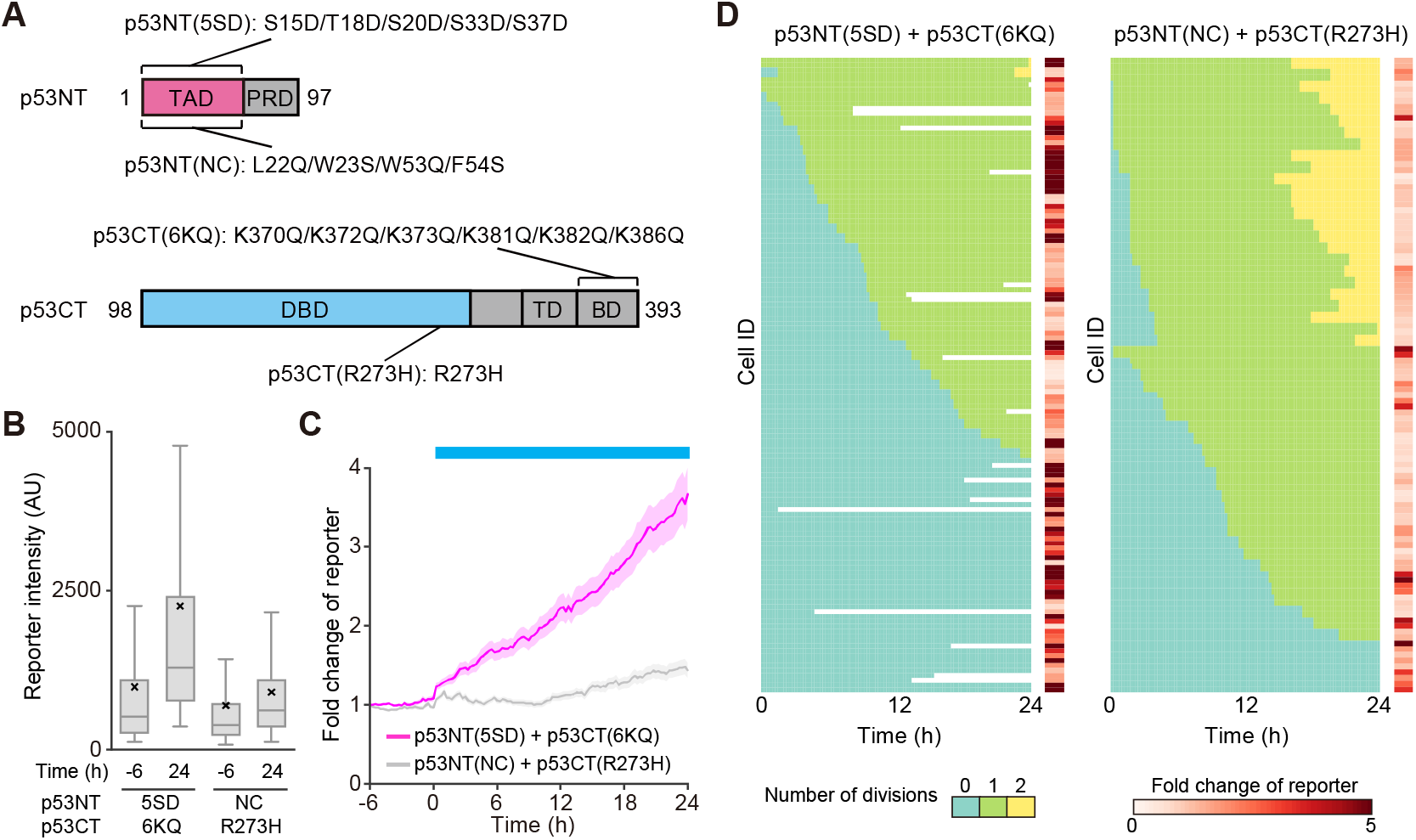
Cell cycle regulation by light-dependent p53 activation. A. Schematic illustration of the p53 domain mutant used in this study. B. Distribution of the fluorescence intensity in transcription reporter at -6 hours and 24 hours from the start of blue light illumination. Horizontal lines and crosses indicate the medians and means of distribution, respectively. Boxes and whiskers include the values between the 25th and 75th percentiles or between the maximum and minimum values excluding outliers, respectively. More than 180 cells were analyzed under each condition. C. Fold changes in the p53 transcription reporter in each cell line. The plot shows the mean ± s.e.m. p53NT(5SD) + p53CT(6KQ), n = 130 cells; p53NT(NC) + p53CT(R273H), n = 110 cells. D. Individual cell division profiles and fold changes in the p53 transcription reporter in Figure 4B. Each row represents the division history of a single cell, with each mitotic event marked by a color transition. Fold changes in the p53 transcription reporter were the ratio of 0 hour to 24 hours if the cell was alive during the observing time window, or the ratio of 0 hour to the value just before death if the cell died during the observation.

We confirmed light-dependent subcellular localization changes in these mutants of Opto-p53 as in Figure 2 (Fig. S3A, B). The expression of only p53NT(5SD) showed foci formation upon blue light (Fig. S3A, upper left). Interestingly, the single expression of p53NT(NC), incapable of binding to the transcriptional coactivator, failed to form foci induced by light (Fig. S3A, upper right), suggesting the involvement of the interaction with transcriptional coactivators such as p300/CBP in condensate formation (Ma *et al*., 2021). The co-expression of actuators with localizers p53CT(6KQ) or p53CT(R273H) promoted light-induced foci formation co-localized with the actuators p53NT(5SD) or p53NT(NC) (Fig. S3A, middle and lower rows). Of note, the number of condensates was reduced in cells expressing p53CT(R273H) compared to p53CT(6KQ) (Fig. S3C), and the size of each condensate was larger (Fig. S3D, E). These data imply that the p53CT bound to the DNA acts as a seed of condensates, thereby increasing the number of foci and decreasing their size.

By using cell lines stably expressing p53NT(5SD) + p53CT(6KQ) (termed 5SD+6KD) and p53NT(NC) + p53CT(R273H) (termed NC+R273H), we examined the p53 transcriptional activity and cell cycle. As we expected, the 5SD+6KQ cells demonstrated significant p53 transcriptional activation after blue light illumination (24 hours), whereas the NC+R273H cells showed only a slight increase (Fig. 4B). In line with the transient expression experiments (Fig. 3B), mScarlet-I fluorescence intensity gradually increased in the 5SD+6KQ cells upon blue light, whereas the NC+R273H cells did not (Fig. 4C). We note that the higher mScarlet-I fluorescence intensity in the basal state (before 6 hours) in 5SD+6KQ cells compared to NC+R273H cells may be due to the slight interaction of p53CT localizer and p53NT actuator under dark conditions.

Finally, we examined the direct relation between p53 transcriptional activity and cell cycle arrest. For this purpose, the 5SD+6KQ and NC+R273H cells were imaged under the blue light condition, and the fluorescence intensity of the p53 transcription reporter and mitotic events were analyzed based on the cell lineage tracking with their time-lapse images. In Figure 4D, the heatmaps, with the time after blue light exposure on the horizontal axis and Cell ID on the vertical axis, indicate color codes representing the number of cell divisions (zero times, once, twice during 24-hour time-lapse imaging). On the right side of heatmaps, the fold changes in p53 transcriptional reporter at the time = 24 hours or the time when cells died are shown in each cell. The cell cycle progression was markedly suppressed in the 5SD+6KQ cells (Fig. 4D, left); about 37% of the cells (48/130 cells) did not show any cell division in the 24-hour period, while the 60% only divided once (78/130 cells). In addition, some cells (18/130 cells) died during observation (white areas in the heat map). In clear contrast, nearly half of the NC+R273H cells (50/110 cells) experienced two cell divisions, and cell death was not observed at all (Fig. 4D, right). Furthermore, the fold change in the p53 transcriptional reporter seemed to be well correlated with the cell fates in these heatmaps; the higher p53 transcriptional activity, the higher the probability of cell cycle arrest and cell death (Fig. 4D).

## Discussion

In this study, we succeeded in developing an optogenetic system, Opto-p53, to activate the p53 signaling pathway in a light-dependent manner. Opto-p53 is a unique system, in which a DBD and TAD derived from an endogenous transcription factor are employed (Fig. 1). Therefore, the design of the light-responsive transcription factor presented in this study provides a general framework for the optogenetic manipulation of endogenous transcription factors.

By using Opto-p53, we observed light-induced transcriptional activation of p53 (Fig. 3A and 3B). Compared with the results of drug treatment to the reporter cell line (Fig. S2B and S2C), the increase in p53 transcription reporter in cells expressing Opto-p53 (approximately 4-fold) was equivalent to that induced by 20 μM etoposide treatment and 2.5 μM nutlin-3a treatment (Figs. 3B and 4C). These drug concentrations were sufficient to induce cell cycle arrest and cell death (Tsuruoka *et* al., 2023; Vassilev *et al*., 2004; Yang *et al*., 2018), indicating that Opto-p53 has the ability to activate p53 to the extent that physiologically affects the cellular phenotype. Indeed, these cells eventually caused cell death and cell cycle arrest (Figs. 3C and 4D). The cells transiently expressing Opto-p53 demonstrated cell death upon blue light (Fig. 3C), while most of the cells stably expressing Opto-p53 showed cell cycle arrest by light (Fig. 4D), implicating that the differences in the phenotypes depend on the amount of expression level of Opto-p53. The remarkable fact is that an endogenous transcription factor-based optogenetic system can reproduce the function of transcription factors. When cells stably expressing Opto-p53 were exposed to blue light, more than half of the cells experienced a single cell division within 24 hours (Fig. 4D). However, most of these cells did not proceed to a second cell division. In contrast, approximately 40% of the cells expressing the negative control fragment underwent two rounds of cell division. This result suggests that cells have the potential to pass through the G2/M checkpoint after p53 activation but not the G1/S checkpoint. This implication is consistent with the classical paradigm that p53 is primarily involved in G1/S checkpoint regulation and, to a lesser extent, in the G2/M checkpoint (Giono and Manfredi, 2006). Another possible explanation is that there exists a time lag between the start of light illumination and the activation of p53, and that only cells entering the M phase before p53 was fully activated underwent cell division.

Transcriptional activation of Opto-p53 monitored by the p53 reporter system exhibited a smaller fold increase than previously reported synthetic transcription factors, which had showed more than a 10-fold increase in transcriptional activation (Gao et al., 2016; Nihongaki et al., 2015; Polstein and Gersbach, 2015; Schneider *et al*., 2021; Wang et al., 2012; Yamada *et al*., 2018). As an actuator, p53TAD itself showed a transcriptional activation comparable to the commonly used transcriptional activation domain derived from VP16, negating the possibility that the transcriptional activity of p53 TAD is insufficient. There are several possible explanations. In previous studies on synthetic transcription factors, TAD-containing fragments are, in principle, excessive compared to the limited number of binding sites of transcription factors in the genome. Meanwhile, in Opto-p53, DBD-containing localizers are expected to interact with numerous p53-responsive elements widely distributed in the genome, thereby showing the apparent difference in the stoichiometry between previously reported synthetic transcription factors and Opto-p53 (Chang *et al*., 2014). In addition, we used HCT116 cell lines possessing the wild-type *TP53* gene in this study. Therefore, we should note the possibility that the endogenous p53 elevates transcriptional activity under basal conditions.

Actuator and localizer fragments of Opto-p53 formed condensate in the nucleus upon blue light illumination (Fig. 2 and Fig. S3). This could be mediated through multiple factors, including the homo-oligomeric activity of CRY2, the interaction of TAD with other transcription-related factors, IDR nature of p53NT, and the ability of tetramer formation in the Opto-p53 localizer. In many cases, the condensation of transcription-related factors is known to increase the transcription efficiency of both endogenous and synthetic transcription factors (Hnisz et al., 2017; Schneider et al., 2021; Wei et al., 2020; Wu *et al*., 2022). On the other hand, it has also been reported that transcriptional activity is repressed depending on the nature of the condensate (Chong *et al*., 2022; Fischer *et al*., 2024). Hence, the relationship between condensate formation and the transcriptional activity of Opto-p53 would require further investigation. There have been reports of optogenetic methods to modulate the properties of condensate, such as optoDroplet and Corelet (Bracha *et al*., 2018; Shin et al., 2017).

Therefore, applying these techniques may more quantitatively reveal the relation between transcriptional activity and condensate properties.

We discuss possible future directions of Opto-p53. One possible direction is to investigate how downstream gene expression is altered by using various mutants of the p53. Endogenous transcription factors are regulated by a lot of context-dependent PTMs (Filtz et al., 2014). p53 is also known to undergo a great variety of PTMs, and over the years, many researchers have devoted their efforts to discovering and assigning roles to each PTM (Wen and Wang, 2022). Based on the design of the light-responsive transcription factors in this study, it would be possible to directly address the role of each PTM in the regulation of transcription factors by introducing a mutation that is not post-translationally modified or a mutation that mimics PTMs. Investigating how different p53 temporal dynamics induce different cellular phenotypes by employing specific light illumination patterns would also be interesting. The advantage of optogenetics, which allows for local activation, could also be applied to studying p53 function in an *in vivo* system. Combining these approaches with Opto-p53 provides deeper insight into regulatory mechanisms of p53 function in the future.

## Materials and Methods

### Plasmids

All plasmids used in this study are summarized in Table S1, along with Benchling links to the plasmid sequences and maps. The oligonucleotides for PCR and DNA sequencing were purchased from FASMAC. The cDNA of human p53 and the expression backbone of the p53 transcription reporter were synthesized by FASMAC. For the p53 transcription reporter vector, the fundamental design followed a previous report (Tsuruoka *et al*., 2023). First, the synthesized expression backbone was transferred to the PiggyBac transposon vector (Yusa et al., 2009). Next, the promoter sequence was replaced with a promoter region derived from the *CDKN1A* gene, which was obtained by PCR using genomic DNA extracted from HCT116 cells as a template by QuickExtract DNA Extraction Solution (Lucigen). The 3xNLS sequence derived from simian virus large T-antigen and mRNA degradation sequence (AU-rich element (AU1): 5′-ATTTATTTATTTATTTATTTA-3′) were subsequently inserted into the reporter gene cassette using DNA fragments obtained by oligo DNA annealing. For the expression vectors of the Opto-p53 fragments, CRY2 and CIBN were obtained from pCAGGS-hCRY2-mCherry (plasmid#178525: Addgene) (Yamamoto et al., 2021) and pCX4neo-CIBN-EGFP-KRasCT (Aoki et al., 2013), respectively. All the expression vectors were constructed by restriction enzyme digestion and ligation or Gibson assembly with NEBuilder HiFi DNA assembly (New England Biolabs).

### Cell culture

HCT116 cells were obtained from the American Type Culture Collection (ATCC; Rockville, MD, USA). Cells were cultured in RPMI 1640 Medium (ATCC modification) (A10491-01: ThermoFisher Scientific) supplemented with 10% fetal bovine serum (FBS; 175012: NICHIREI). All cells were maintained in a humidified atmosphere of 5% CO_2_ at 37°C.

### Transfection

Cells were plated on a collagen-coated 4-well glass-bottom dish (The Greiner Bio-One) for 1 day and transfected with 0.5 μg of each plasmid by using Polyethyleneimine ‘Max’ MW 40,000 (Polyscience Inc., Warrington, PA, USA). Time-lapse imaging was started 1 day after transfection.

### Establishment of stable cell lines

Cells were transfected with PiggyBac donor vectors and PiggyBac transposase-expressing vectors at a ratio of 3:1 (Yusa *et al*., 2011). Nucleofector IIb (Lonza, Basel) electroporation system was used for transfection according to the manufacturers’ instructions (D-032 program) with a house-made DNA- and cell-suspension solution (4 mM KCl, 10 mM MgCl_2_, 107 mM Na_2_HPO_4_, 13 mM NaH_2_PO_4_, 11 mM HEPES pH 7.75) (Yamamoto et al., 2021). One day after transfections, cells were treated with 1 μg/mL puromycin (InvivoGen, San Diego, CA), 10 μg/mL blasticidin S (InvivoGen), or G418 (InvivoGen) for drug selection. Time-lapse imaging for stable cell lines was started 2 days after seeding on a collagen-coated 4-well glass-bottom dish. For Figures S2B and S2C, the stable cell line harboring p53 transcription reporter was seeded to a 96-well flat-bottom microplate (The Greiner Bio-One).

### Live-cell fluorescence imaging

Epifluorescence inverted microscopes (IX83; Olympus, Tokyo) were used for live-cell imaging. For spinning-disk confocal microscopy (Figures 2, S1, and S3), the microscope was equipped with an sCMOS camera (ORCA Fusion BT; Hamamatsu Photonics), oil-immersion objective lens (UPLXAPO 60X, NA = 1.42, WD = 0.15 mm; Olympus), and a spinning-disk confocal unit (CSU-W1; Yokogawa Electric Corporation). The excitation lasers and fluorescence filters used were as follows: Excitation laser, 488 nm for mNeonGreen, 561 nm for mCherry, and 640 nm for miRFP703; excitation dichroic mirror, DM405/488/561/640 for mNeonGreen, mCherry, and miRFP703; emission filters, 525/50 for mNeonGreen, 617/73 for mCherry, and 685/40 for miRFP703 (Yokogawa Electric). For wide-field fluorescence microscopy (Figures 3, 4, and S2), the microscope was equipped with a Prime sCMOS camera (Photometrix) and dry objective lens (UPLXAPO 40X). The illumination settings and fluorescence filters were as follows: Excitation wavelength, 475/28 for mNeonGreen, 580/20 for mScarlet-I, and 632/22 for miRFP703; excitation dichroic mirror, FF409/493/573/652/759-Di01-25.8×37.8 for all fluorescent proteins; emission filters, 520/28 for mNeonGreen, 641/75 for mScarlet-I, and 664/long pass (Semrock). The illumination light source was a Spectra X light engine (Lumencor). For the illumination of the blue light, blue LED light (450 nm) (LED-41VIS450; OptoCode Corp., Japan) was continuously illuminated from the top of the stage.

For Figures S2B and C, ImageXpress Micro XLS (Molecular Devices) was used for live-cell imaging. The microscope was equipped with an sCMOS camera (Zyla 5.5; Andor) and dry objective lens (Plan Fluor 10X, NA = 0.30, WD = 16 mm; Nikon). The illumination settings and fluorescence filters were as follows: Excitation wavelength, 562/40 for mScarlet-I; excitation dichroic mirror, 350-585 (R)/601-950 (T) for mScarlet-I; emission filters, 624/40 for mScarlet-I (Semrock). The illumination light source was a SOLA SE2 (Lumencor).

### Image Analysis

All imaging data were analyzed using Fiji/ImageJ (Schindelin et al., 2012). First, the background was subtracted for all images by the rolling-ball method, and then stacked images were created. Next, LIMTracker, a Fiji tracking plugin, was used to detect nuclear regions and track cell trajectories, and the temporal changes in fluorescence profiles were quantified for each cell trace (Aragaki et al., 2022). Nuclear regions were detected based on the H1-mCherry signal (Figures 2, S1, and S2) or p53 transcriptional reporter signal (mScarlet-I-3xNLS; Figures 3, 4, and S2).

For the data of Figures 4A, S2B, and S2C, the Fiji plugin StarDist was used to detect the nuclei of all cells in the images at each time point, followed by quantification of the fluorescent signal in the nuclei. All the quantified data were analyzed and visualized using Python with seaborn and matplotlib.

For Figures S3C∼E, cells with a CV value larger than 0.4 were selected in order to analyze only cells with foci. The foci areas were detected as follows.

1. Create cropped images of cells from the frame 30 minutes after the light illumination.
2. Apply Top Hat processing to the cropped images.
3. Binarize the cropped images by Otsu’s method.
4. Detect the number and size of foci areas based on the binarized images.

## Acknowledgments

We thank all members of the Aoki Laboratory for their helpful discussions and assistance. We thank E. Ebine-Sato, K. Onoda, M. Hirao, and T. Uesugi for their technical support and the other members of the Aoki laboratory for their helpful discussions.

