## Supplementary Materials for "Opto-p53: A Light-Controllable p53 Signaling for Regulating p53-Dependent Cell Fate"

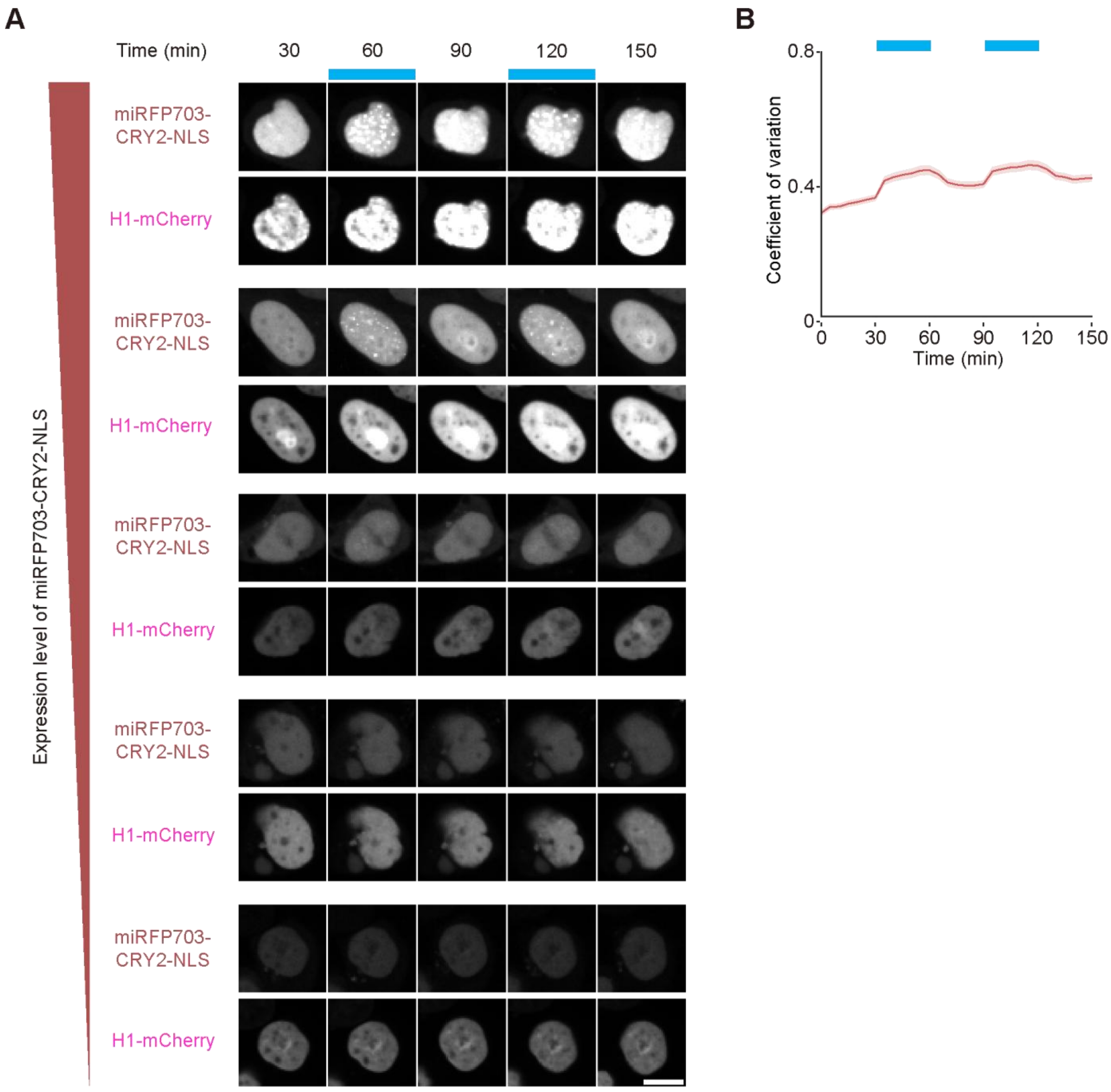


**Figure S1. CRY2 alone forms condensates upon blue light illumination only at high expression levels.**

A. Light-dependent changes in subcellular localization of miRFP703-CRY-NLS fragments with different expression levels. The blue boxes indicate the time points at which blue light illumination was applied. H1-mCherry is a nuclear marker. Scale bar, 10 µm.

B. Temporal changes in the coefficient of variation of nuclear miRFP703 fluorescence intensity in cells expressing miRFP703-CRY-NLS. The plot shows the mean ± s.e.m. n = 91 cells.

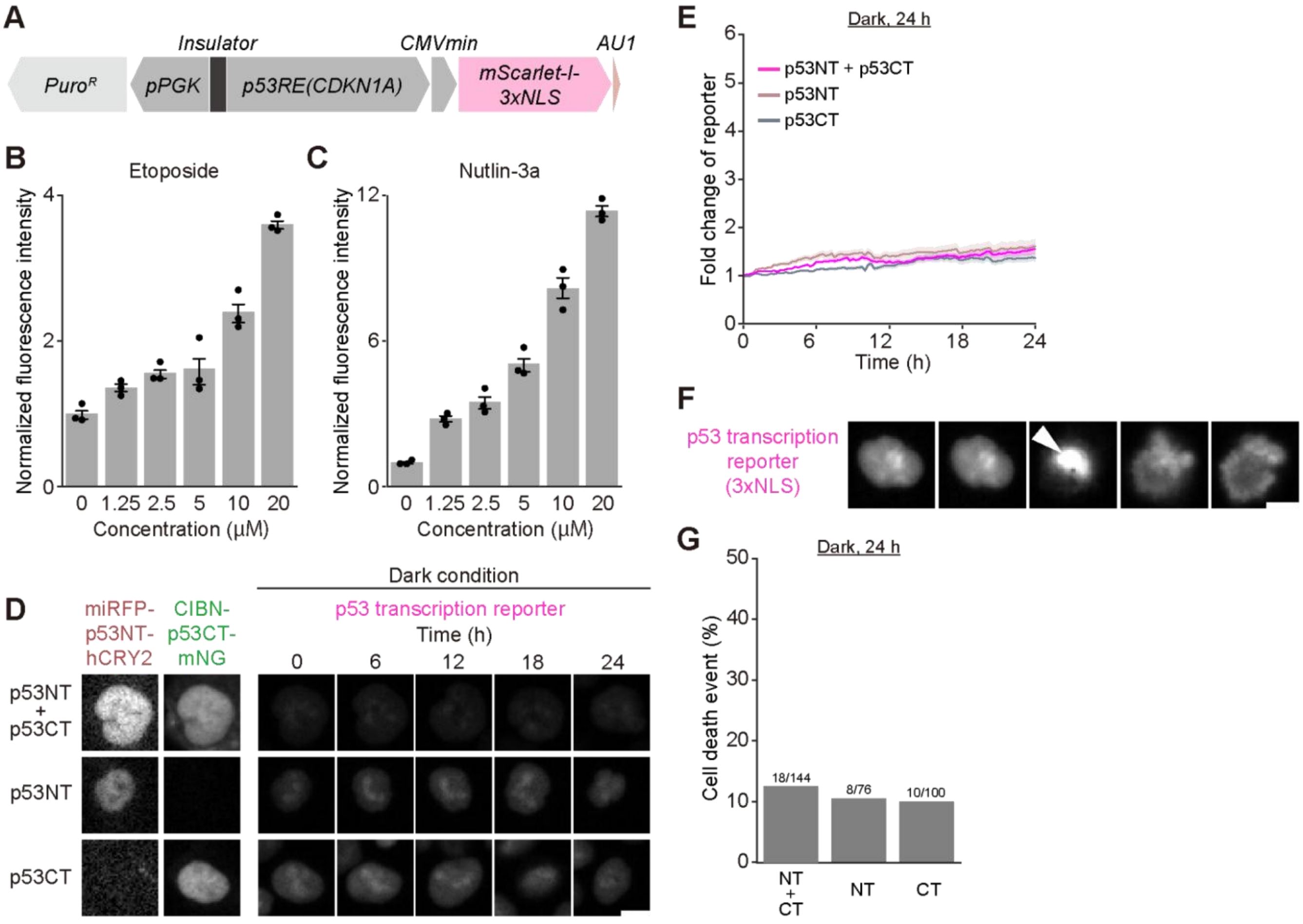


**Figure S2. Functional characterization of the p53 transcriptional reporter and behavior of Opto-p53 under dark conditions.**

A. Schematic representation of the reporter system used in this study. Puro$^R$, Puromycin resistance gene; pPGK, phosphoglycerate kinase promoter; CMVmin, CMV minimal promoter; AU1, AU-rich element.

B. Dose-response of the stable cell line harboring p53 transcription reporter 24 hours after treatment with the indicated concentration of etoposide.

C. Dose-response of the stable cell line harboring p53 transcription reporter 24 hours after treatment with the indicated concentration of nutlin-3a.

D. Expression patterns of Opto-p53 actuator or localizer and changes in p53 transcription reporter under dark conditions. Scale bar, 10 μm.

E. Fold changes in the p53 transcription reporter under dark conditions. The plot shows the mean ± s.e.m. p53NT + p53CT, n = 144 cells; p53NT, n = 76 cells; p53CT, n = 100 cells.

F. Morphological changes in cell death. White arrowheads indicate chromatin compaction associated with cell death. Cell images were obtained every 15 minutes.

G. Quantification of cell death in each condition. The numbers on each grey bar indicate the number of dead cells/total cells.

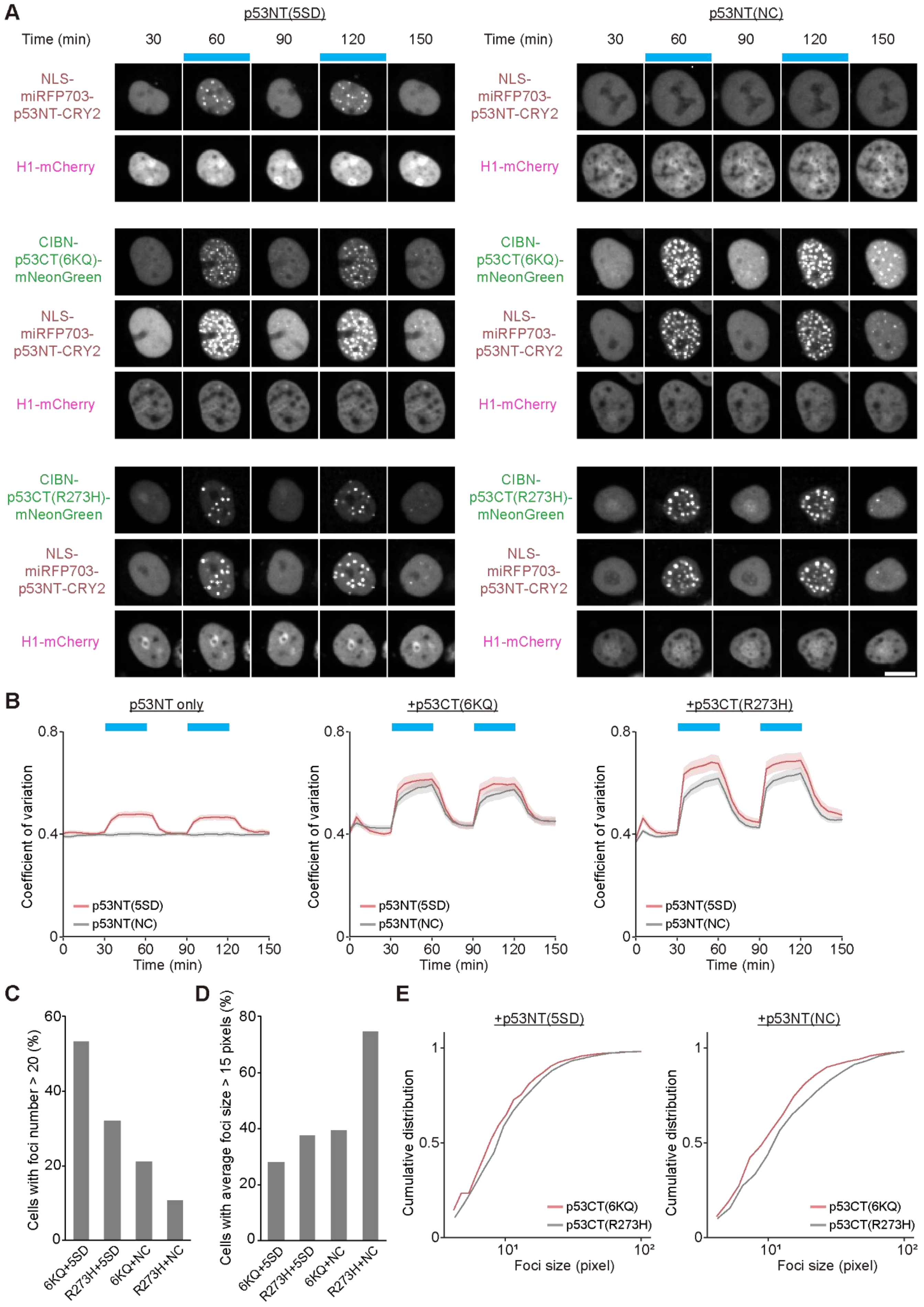

A
p53NT(5SD)
p53NT(NC)
Time (min)
30
60
90
120
150
NLS-miRFP703-p53NT-CRY2
H1-mCherry
CIBN-p53CT(6KQ)-mNeonGreen
CIBN-p53CT(R273H)-mNeonGreen
B
p53NT only
+p53CT(6KQ)
+p53CT(R273H)
Coefficient of variation
p53NT(5SD)
p53NT(NC)
Time (min)
C
Cells with foci number > 20 (%)
6KQ+5SD
R273H+5SD
6KQ+NC
R273H+NC
D
Cells with average foci size > 15 pixels (%)
E
+p53NT(5SD)
+p53NT(NC)
Cumulative distribution
p53CT(6KQ)
p53CT(R273H)
Foci size (pixel)

**Figure S3. Subcellular localization changes in Opto-p53 fragments with mutations in the p53 domain.**

A. Light-dependent changes in subcellular localization of each Opto-p53 actuator or localizer with different mutations in the p53 fragment. The blue boxes indicate the time points at which blue light illumination was applied. H1-mCherry is a nuclear marker. Scale bar, 10 µm. Upper panels, p53NT alone; middle panels, p53CT(6KQ) mutant + p53NT; lower panels, p53CT(R273H) mutant.
B. Temporal changes in the coefficient of variation of nuclear miRFP703 fluorescence intensity for each condition as shown in Figure S2A. Left panel, p53NT alone; middle panel, p53CT(6KQ) mutant + p53NT; right panel, p53CT(R273H) mutant. The plot shows the mean ± s.e.m. p53NT(5SD), n = 127 cells; p53NT(NC), n = 120 cells; p53CT(6KQ) + p53NT(5SD), n = 122 cells; p53CT(6KQ) + p53NT(NC), n = 151 cells; p53CT(R273H) + p53NT(5SD), n = 118 cells; p53CT(R273H) + p53NT(NC), n = 159 cells.
C. Cell population with the foci number > 20 in each co-expression condition.
D. Cell population with the average foci size > 15 pixels in each condition.
E. Cumulative distribution of foci size in all detected foci. Left panel, p53CT(6KQ) mutant + p53NT; right panel, p53CT(R273H) mutant. *p*-value: $3.0 \times 10^{-6}$ for p53CT(6KQ) + p53NT(5SD) vs p53CT(6KQ) + p53NT(NC), $5.1 \times 10^{-8}$ for p53CT(R273H) + p53NT(5SD) vs p53CT(R273H) + p53NT(NC). *p*-values were calculated by a Mann-Whitney U test.

**Table S1**

| Plasmid name | Figures | Source or reference | Sequence |
|---|---|---|---|
| pCSIIbleo-H1-mCherry | 2, 3, S1, S3 | This study | https://benchling.com/s/seq-8w1sVGlEYvw4uyexxMKw?m=slm-gKtz0EOAkyiXOqhoKjtM |
| pPBbsr2-miRFP703-dSal-p53NT(1-97)-S15D-hCRY2-NLS | 2, 3, S2 | This study | https://benchling.com/s/seq-iaymBMZnxtC3UG3iM9Kj?m=slm-Ki8MNcdXMSnSiICJc5WF |
| pPBbsr2-miRFP703-dSal-VP16minADx3-hCRY2-NLS | 2, 3 | This study | https://benchling.com/s/seq-Z27cmjXf0ZOWdL9Q0ZE7?m=slm-tU3F0ok9Pb2DYIy0k3ZJ |
| pCAGGS-CIBN-p53CT(98-393)-mNeonGreen | 2, 3, S2 | This study | https://benchling.com/s/seq-SmXmLCJPBWd2Ezilwum3?m=slm-V9iSpDwed2bFAJ3BnxFV |
| pPB-p53RE(CDKN1A)-CMVmin-mScarlet-I-NLSx3-AU1-PGKpuro | 3, 4, S2 | This study | https://benchling.com/s/seq-lv1G3m5olUoQKYBtBMca?m=slm-CbOooCTkrhJDY4fPlsMi |
| pPBbsr2-NLSx3-miRFP703-dSal-p53NT(1-97)-5SD-lin-hCRY2 | 4 | This study | https://benchling.com/s/seq-fZiB5aeNnaWCmscug3kT?m=slm-gPuQW5SitzBdnclEQHhi |
| pPBbsr2-NLSx3-miRFP703-dSal-p53NT(1-97)-L22Q/W23S/W53Q/F54S-lin-hCRY2 | 4 | This study | https://benchling.com/s/seq-m6oqqZOadMHkaQ9eZi2E?m=slm-bQNOzruUFTAGNoZLV7Hl |
| pPBneo-CIBN-lin-p53CT(98-393)-6KQ-mNeonGreen-NLSx3 | 4 | This study | https://benchling.com/s/seq-xlq0aODx512OPaNJg2Kh?m=slm-SrXB4VYMFipwHP2l6PkD |
| pPBneo-CIBN-lin-p53CT(98-393)-R273H-mNeonGreen-NLSx3 | 4 | This study | https://benchling.com/s/seq-R0dnqxz6dlOdlaldc5Yb?m=slm-UYRM4hZLBiayM7JI2GFL |
| pPBbsr2-miRFP703-dSal-lin-hCRY2-NLS | S1 | This study | https://benchling.com/s/seq-lMmo585RZDRqqFKmtzxf?m=slm-pwb8jrlqveI9pc8yBtYS |
| pPBbsr2-SuperNLS-miRFP703-dSal-p53NT(1-97)-5SD-lin-hCRY2 | S3 | This study | https://benchling.com/s/seq-WMn7te2KkdVCZ6pTtptT?m=slm-qHednjcL97CTyZZk3pWC |
| pPBbsr2-SuperNLS-miRFP703-dSal-p53NT(1-97)-L22Q/W23S/W53Q/F54S-lin-hCRY2 | S3 | This study | https://benchling.com/s/seq-1eUakm7jJFRSaz6FaAt3?m=slm-ewWwpMXkDH3YEo2u0WmH |
| pPBneo-CIBN-lin-p53CT(98-393)-6KQ-mNeonGreen-SuperNLS | S3 | This study | https://benchling.com/s/seq-T3aCEpiu08SmIh1NcjCh?m=slm-xlLyqvUYU2a9LxdKieqJ |

| | | | |
|---|---|---|---|
| pPBneo-CIBN-lin-p53CT(98-393)-R273H-mNeonGreen-SuperNLS | S3 | This study | https://benchling.com/s/seq-S6yezfzJ0OD1iqbU2U0a?m=slm-ZpfYeI4bB5vzkFqNAyaw |

**Table S1. Plasmid list used in this study.**

**Movie S1**
HCT116 cells transiently expressing the p53NT actuator were cultured on a 4-well glass-bottom dish. Time-lapse imaging was performed using a spinning disk confocal microscope. The cells were repeatedly illuminated with blue light at 30 min intervals. Images were acquired every 5 min. Total imaging time = 150 min.

**Movie S2**
HCT116 cells harboring a p53 transcriptional reporter were cultured on a 4-well glass-bottom dish and transiently expressing Opto-p53. Time-lapse imaging was performed using a wide-field microscope. The cells were continuously illuminated with blue light at 24 hours. Images were acquired every 15 min. Total imaging time = 24 hours.
